# Comparative proteomics uncovers distinct biomarkers, protein networks and defense responses in tomato during beneficial and pathogenic microbial interactions

**DOI:** 10.1101/2025.01.23.634544

**Authors:** Dhananjaya Pratap Singh, Sudarshan Maurya, Suresh Reddy Yerasu, Ratna Prabha, Renu S, Birinchi K. Sarma, Lovkush Satnami, Nagendra Rai

**Affiliations:** ICAR-Indian Institute of Vegetable Research, Shahanshahpur, Varanasi 201305, India; ICAR-Indian Agricultural Research Institute, Agricultural Knowledge Management Unit, 110012 New Delhi, India; Indian Council of Agricultural Research, Krishi Bhawan, 110001 New Delhi, India; Department of Mycology and Plant Pathology, Institute of Agricultural Sciences, Banaras Hindu University, Varanasi 221005 India

**Author notes:** Corresponding author: Dhananjaya Pratap Singh, Principal Scientist (Biotechnology).

**Keywords:** Proteomics, LC-MS, Plant-microbe interactions, Tomato, *Bacillus subtilis*, *Alternaria solani*, Biomarkers

## Abstract

The molecular mechanisms underlying plant responses to beneficial and pathogenic microbial interactions has been uncovered in tomato at the proteomic level. Our study employed LC-MS-based proteomics to investigate differential protein regulation in tomato plants during interactions with beneficial bacteria (*Bacillus* subtilis BV7) versus pathogenic fungus (*Alternaria solani*). Comparative analysis revealed distinct protein signatures characterizing each interaction: 232 unique proteins in BV7-treated plants versus 96 in pathogen-infected plants, with 54 proteins shared between treatments. BV7 inoculation enhanced proteins involved in photosynthesis and primary metabolism, with PSI-K emerging as the top biomarker (score 2.53), while pathogen infection triggered focused defense responses with Cytochrome b559 (score 2.38) as the key biomarker. Metabolic pathway analysis demonstrated that BV7 uniquely enhanced vitamin metabolism (thiamine, riboflavin, folate) and energy production pathways, while pathogen infection activated defense-related phenylpropanoid biosynthesis. Analysis of defense enzymes showed pathogen infection induced highest activities of PAL (39.4±0.46 U h^-1^ g^-1^ fw), SOD (29.9±0.94 U mg^-1^ protein), POD (3.18±0.07 μg g^-1^ fw min^-1^), APx (15.58±0.23 U mg^-1^ protein), and GPx (71.51±0.7 U mg^-1^ protein), while BV7 maintained moderate enzyme levels, suggesting balanced growth-defense responses. The shared proteins between treatments indicate a common molecular framework potentially contributing to induced systemic resistance. These findings provide novel insights into plant-microbe interactions at the molecular level, identifying specific protein biomarkers and metabolic pathways that could be targeted for enhancing crop productivity and disease resistance. The results have significant implications for developing biological control strategies and improving sustainable agricultural practices.

## 1. Introduction

The intricate interactions between plants and microorganisms play a pivotal role in shaping plant health, productivity, and overall ecosystem dynamics. Plant-microbe interactions can be broadly categorized as beneficial, neutral, or pathogenic, each eliciting distinct molecular responses within the host plant (Jones et al, 2019). Understanding the molecular mechanisms underlying diverse microbial interactions with plants is crucial for developing sustainable agricultural practices and enhancing crop resilience in the face of biotic and abiotic stresses (Agrahari et al, 2020). Tomato (*Solanum lycopersicum*), a globally important vegetable crop, serves as an excellent model system for studying plant-microbe interactions due to its economic significance, genetic tractability, and susceptibility to various pathogens (Krishna et al, 2022, Liu et al, 2022). The plant’s responses to beneficial and pathogenic microbes provide valuable insights into the complex molecular networks that govern plant immunity, growth promotion, and stress tolerance.

Beneficial gram-positive bacterium *Bacillus subtilis* has widely recognized as a plant growth-promoting rhizobacterium (PGPR) that colonizes plant rhizosphere and confers multifaceted benefits to the host plant, including enhanced nutrient uptake, phytohormone production, and induced systemic resistance (ISR) against pathogens (Vejan et al, 2016). The mechanisms by which *B. subtilis* promotes plant growth and health involve complex signaling cascades and metabolic reprogramming within the host plant (Hashem et al, 2019). On the other hand, *Alternaria solani* causes a devastating early blight disease in tomato and significantly impacts production of this crop worldwide. This necrotrophic fungus induces cell death and tissue necrosis, leading to substantial yield losses. The interaction between tomato and *A. solani* triggers a distinct set of biochemical and molecular mechanisms leading to defense responses, including the activation of pathogenesis-related (PR) proteins, phytoalexin production, and hormonal signaling pathways (Pandey et al, 2023). The contrasting nature of beneficial and pathogenic interactions provides a unique opportunity to elucidate the molecular signatures that differentiate plant responses to diverse microbial stimuli. While previous studies have focused on individual aspects of these interactions, a comprehensive understanding of the global proteomic changes and associated pathway shifts remains elusive.

Recent advances in high-throughput proteomics technologies have revolutionized our ability to study complex biological systems at unprecedented depth and resolution (Yan et al, 2022). These approaches enable the identification and quantification of thousands of proteins simultaneously, offering a holistic view of cellular processes and regulatory networks (Hu et al, 2015). To provide a comprehensive understanding of the differential responses of tomato to beneficial and pathogenic interactions, we have applied state-of-the-art proteomics techniques to study *B. subtilis* and *A. solani* interaction at proteomic level and unravel the intricate molecular landscape that defines contrasting relationships. The objective of this study was to decipher the differential proteomic signatures and pathway shifts in tomato plants in response to colonization by the beneficial *B. subtilis* and infection by the pathogenic *A. solani*. By comparing distinct proteomic profiles, we have identified key biomarker proteins, metabolic pathways, and regulatory networks that are specifically modulated in each interaction. This comparative approach will provide valuable insights into the molecular mechanisms underlying plant growth promotion, induced resistance, and pathogen defense. Our findings not only advance fundamental knowledge of plant-microbe interactions at the proteomic level but also pave the way for innovative strategies to enhance crop resilience and productivity in the face of diverse microbial challenges. The insights gained from the findings may help in targeting important pathways for manipulation through microbial inoculations for reducing chemical farm inputs and promoting environment-friendly crop protection methods.

## 2. Material and Methods

### 2.1. Beneficial microbe, pathogen and plant treatments

Seeds of the tomato plants (*S. lycopersicum* var. Kashi Aman) were sown in pots (20x20x14cm) filled with sterilized field soil, cocopeat, perlite and vermiculite in a ratio of 3:1:1:1; w/w. Potted seeds were kept in greenhouse conditions (Singh et al, 2023, 2024). After emergence, seedlings were allowed to grow and develop at 25°C/20°C (day/night; D/N) thermal condition and 14h/10h (D/N) photoperiod at a relative humidity of 70%. An artificial light source illuminating 600 μmol m^−2^ s^−1^ of light was used to keep plants illuminated. Uniformly growing 3 weeks old seedlings having similar height and growth were used for the bacterial inoculation treatment. The beneficial bacterium, *Bacillus subtilis* (BV7) was grown in LB broth media (tryptone 10g, yeast extract 5g, NaCl 10g and distilled water make up to 1L) at 37°C. The microbial inoculant using *B. subtilis* BV7 was prepared using the spores of the bacterium (2.7x10^8^ mL^−1^ colony-forming units (CFU) in which the pre-germinated tomato seedlings were dipped for 15 min and then transplanted to the experimental pots. Separate sets of seedlings were transplanted in separate pots having control and bacteria-inoculated plants were allowed to grow under similar growth conditions till 60 days. One set of control plants was inoculated with *A. solani* culture (2.1×10^4^ conidia mL^−1^) on 61^th^ day at their lower leaves. Altogether, three sets of plants were available: control (non-treated), *B. subtilis* BV7 inoculated and *A. solani* infected plants. Plants were maintained at 86-90% relative humidity to facilitate disease progression and sampling was done after 5 days of pathogen inoculation. The disease incidence (%) was noted at the time of sampling. Each treatment was replicated three times.

### 2.2 Enzyme extraction and antioxidant activity

Enzyme extracts were prepared from the same leaf samples as were used for the extraction of global plant proteome. Freshly harvested leaf tissues (0.5 g) from control, BV7-inoculated and pathogen-infected plants were homogenized separately in extraction buffer containing 10 ml sodium phosphate buffer (0.1M), 0.2 mL mercaptoethanol and 1.0 g of polyvinyl pyrrolidone (PVPP). The extracted suspension was kept for 1 h after centrifugation (12000 rpm, 30 min, 4°C) and precipitated with 5 volumes of ammonium acetate (0.1M) in methanol. The enzyme (protein) mixture was stored 1h and then centrifuged (10000 rpm, 10min) to obtain pellets that were finally re-suspended in extraction buffer (3ml). Protein content was determined (Lowry et al., 1951). Assays for the analysis of antioxidant activity related enzymes were performed for phenylalanine ammonia-lyase (PAL), peroxidase (POD), superoxide dismutase (SOD), ascorbate peroxidase (APx) and guaiacol peroxidase (GPx) as per the methods described in Singh et al (2020) and Patani et al, (2023) with minor and conditionally suitable modifications. All the experiments were performed in 3 replications and the data were subjected to the test of statistical significance using SPSS 16.0.

### 2.3. Proteome extraction from plant leaves

Global proteome extraction was performed using the phenol extraction method (Faurobert et al, 2007). The procedure for sample preparation included grinding of 1g of fresh leaf tissues kept in frozen liquid nitrogen in a pre-cooled pestle-mortar. The sample was the suspended in ice-cooled methanol for 1h to remove pigments and phenolic contaminants that interfere with the total proteome content. After centrifugation in cold condition, the residue was re-suspended in 3ml extraction buffer (500mM Tris HCl, 50 mM EDTA, 700 mM sucrose, 100 mM KCl, pH 8.0). Mercaptoethanol (2%) and phenylmethylsulphonyl fluoride (PMSF) (1mM) were added before re-suspension. The mixture was the vortexed and incubated on ice for 10 minutes with shaking. An equal volume of Tris-buffered phenol (phenol + tris-HCl saturated buffer, pH 6.6/7.9) was added. After 10 minutes of incubation, the solution was centrifuged at 12000*g*. The phenolic phase was removed thrice. Extracted protein was precipitated using 0.1M ammonium acetate in cold methanol. After centrifugation, the resulting pellet was stored at -20°C. Protein concentration was determined using Bradford protein assay (Bradford 1976). The precipitated protein was dissolved in G-buffer (guanidine hydrochloride) and the protein solution was filtered through a 0.22µM syringe filter prior to proteomic data generation.

### 2.4 Trypsin digestion and tandem mass tag (TMT) labeling

Protein samples (100µg) were combined with 200µL of 50 mM NH_4_HCO_3_ and digested with trypsin following López-Ferrer et al, (2008). Samples were treated with 5 mM dithiothreitol (DTT) and incubated for 1h at 55°C, followed by the addition of 11 mM iodoacetamide and a 15-minute incubation in dark at room temperature. Urea concentration was reduced to <2M. Trypsin was added at a 1:50 (trypsin:protein) ratio, and samples were digested overnight at 37°C. A second enzymatic hydrolysis was performed with trypsin at 1:100 (trypsin:protein) for 4h. The digested peptides were desalted, freeze-dried, and then dissolved in 0.5M triethylammonium bicarbonate. Finally, the peptides were labeled using a tandem mass tag (TMG) kit according to the manufacturer’s instructions.

### 2.5 LC-MS analysis

Peptide separation was performed using reverse phase nano-liquid chromatography on a Thermo Scientific Easy nLC 1200 instrument (Thermo scientific). The column used was a PepMap RSLC C18 2µm, 75µm x 50cm, maintained at 60°C. A 2µg peptide sample was loaded onto the column. For separation, the solvents were, A: 98% water, 2% acetonitrile, 0.1% formic acid, and B: 20% water, 80% acetonitrile, 0.1% formic acid with a flow rate constant at 300nL min^-1^. The gradient profile was 1. 0-2 min: 5% to 10% solvent B, 2. 2-104 min: 10% to 45% solvent B, 3. 104-105 min: 45% to 90% solvent B, 4. 105-115 min: hold at 90% solvent B, 5. 115-116 min: decrease to 5% solvent B and 6. 116-118 min: hold at 5% solvent B. All the solvents used were LC-MS grade (Fischer Scientific). Internal calibration utilized a lock mass of 445.12003 Da.

MS data was generated using a Q Exactive Orbitrap mass spectrometer (Thermo Scientific) coupled to an Easy-nLC 1200 liquid chromatograph. The run time was 0 to 120 min in positive polarity with a default charge state of 2.0. For MS1, settings included 1 microscan, 70000 resolution, 3e6 AGC, 60ms maximum IT time, and a 350 to 2000 m/z scan range. MS2 settings were 17500 resolution, 200 to 2000 m/z full scan range, 1e5 AGC, 12ms ion transfer, 100 m/z fixed first mass, and 27 normalized collision energy. Data dependent analysis (DDA) settings included 1.00e2 AGC target max, 50.0 S dynamic exclusion time, and unassigned charge exclusion.

### 2.6 Database search

The annotation of proteins in the samples was performed using Proteome Discoverer 2.5 in conjunction with the UniProt protein database of *Solanum lycopersicum* (UniProt). Validation parameters based on q-values included peptide length 6 minimum, 144 amino acid residues maximum, precursor mass tolerance 10 ppm, secondary fragment ion mass error tolerance 0.02 Da and restriction method trypsin/p with maximum allowed missed cleavage up to 2. The false discovery rate (FDR) was restricted to 0.01 for both protein and peptide-spectrum match (PSM) identification.

### 2.7 Functional annotation and enrichment analysis

GO based annotation was done using UniProt-GOA database (http://www.ebi.ac.uk/GOA). Protein IDs were converted to UniProt IDs and then GO IDs were matched to retrieve corresponding information from the UniProt-GOA database. InterProScan was used to predict the GO function of the protein if there was no protein information query on the UniProt-GOA database. Venny 2.1 (Venny 2.1.0 (csic.es) was used to generate a Venn diagram to classify unique proteins in the samples. MetaboAnalyst 6.0 and SIMCA 18 (Sartorius) was implied to perform univariate (volcano plot) analysis and SS Plot was used for data visualization. Data was normalized taking median and log transformation values and Pareto scaling prior to multivariate analysis and preparation of correlation heatmap. Panther classification system (pantherdb.org) database was used to classify GO-Slim biological and molecular functions and categorize protein class. InterPro (a resource for functional analysis of protein sequence family classification, prediction of domains and special sites) database was used to analyze the functional domains of proteins.

The Kyoto Encyclopedia of Genes and Genomes (KEGG) (KEGG: Kyoto Encyclopedia of Genes and Genomes) pathway database of *S. lycopersicum* (*sly*) was used to annotate protein pathways. Pathway mapping of proteins was facilitated by an interactive pathway explorer tool iPATH3.0 (iPath 3: interactive Pathways Explorer (embl.de). The functional protein association network web-tool STRING (STRING: functional protein association networks (string-db.org) was used to perform protein-protein interaction (PPI) networks of the up- and down-regulated proteins identified in the samples. ROC curve analysis module of MetaboAnalyst 6.0 was used for the identification of protein biomarkers. A *t*-test was employed to test the enrichment of the differentially expressed proteins against all identified proteins. The GO/KEGG/protein domains with a corrected *p*-value <0.05 were considered significant.

## 3. Results

The study evaluated how the protein composition changed in the individual set of plant leaves in control (non-treated), beneficial bacteria inoculation and pathogen (*A. solani*) infection condition and analyzed comparative proteome profile to evaluate differential impact of beneficial and pathogenic interactions. We evaluated the activity of defense related antioxidant enzymes PAL, POD, SOD, APx and GPx in the plant leaves for the assessment of biochemical impact of beneficial microbe inoculation and pathogen infection at enzymatic level (Fig 1). Comprehensive biochemical analyses of plant leaf under different treatment conditions, including control (C), beneficial *Bacillus* BV7 (B), and pathogenic *A. solani* (P) treatments were presented in Fig 1. At the time of the sampling for proteome extraction, infected plants showed a disease incidence of 72.4%±1.2 showing high degree of pathogen infection (Fig 1a). Determination of total protein content revealed that *Bacillus* BV7 treatment induced highest protein levels (3.13±0.21mg g^-1^ fw) compared to both the control and pathogen-infected samples, while control and pathogen treatments showed similar, relatively lower protein contents (Fig 1b). The analysis of the activities of several defense enzymes across these treatments showed varying response. PAL activity demonstrated a progressive increase from control to BV7 to pathogen treatments, with the pathogen treatment showing the highest activity (39.4±0.46 U h^-1^ g^-1^ fw) (Fig 1c). Similarly, SOD activity exhibited a stepwise increase across treatments, with control showing the lowest activity, followed by intermediate activity in BV7-treated samples, and the highest activity in pathogen-infected tissues (29.9±0.94 U mg^-1^ protein) (Fig 1d). Regular POD activity followed a similar trend, with control samples showing minimal activity, BV7 treatment inducing moderate activity, and pathogen infection resulting in the highest activity levels (3.18±0.07 μg g^-1^ fw min^-1^) (Fig 1e).

**Figure 1:**
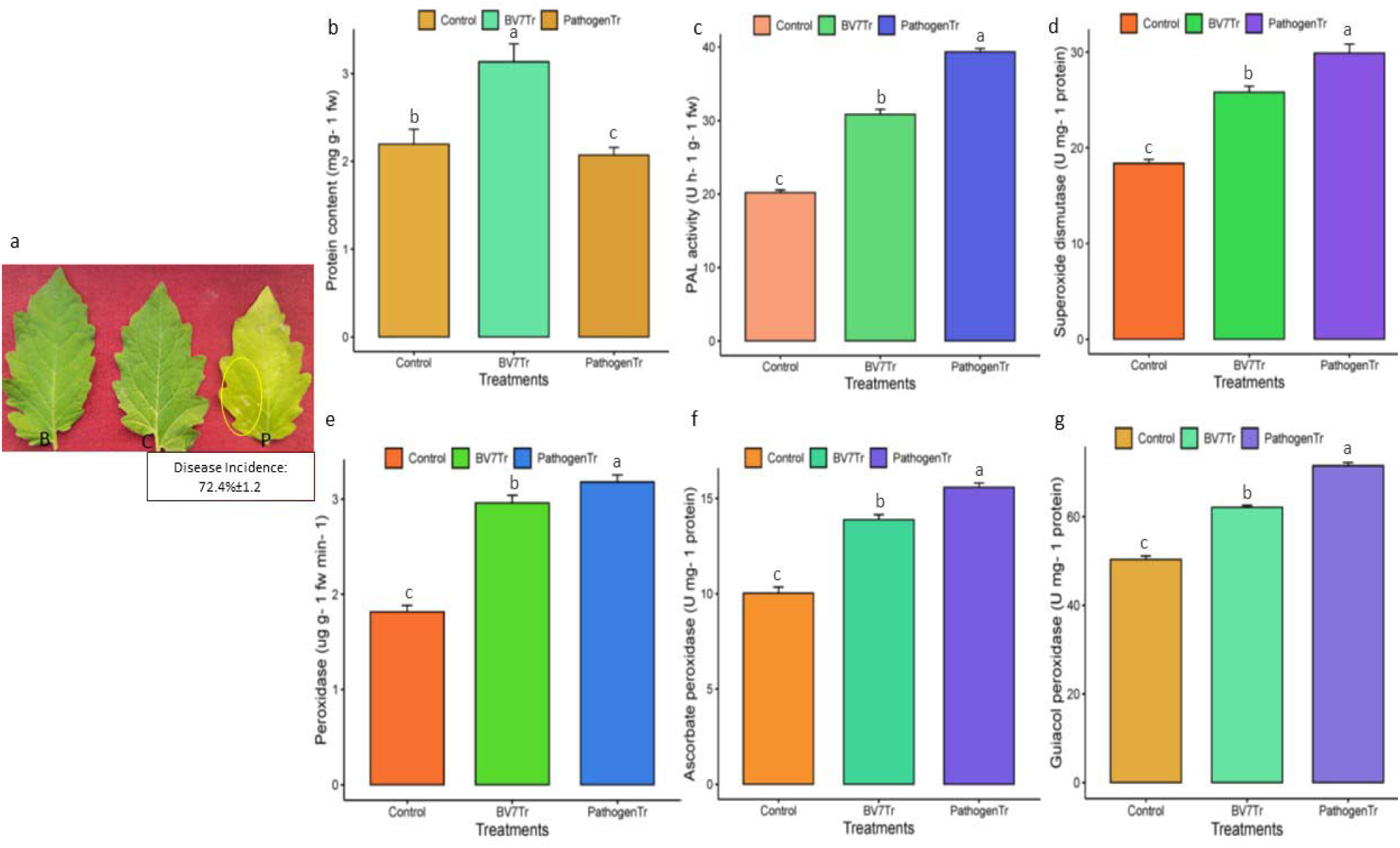
Biochemical and disease responses in plant leaves under different treatment conditions including a. inoculation with the beneficial *Bacillus* BV7 (B), infection with the pathogen *A. solani* (P) and Control (C). The disease incidence was reported to be 72.4%±1.2. Quantitative analyses of b. total protein content, and various biochemical markers including enzymes c. phenylalanine ammonia-lyase (PAL) activity, d. superoxide dismutase (SOD) activity, e. peroxidase activity, f. ascorbate peroxidase activity, and g. guaiacol peroxidase activity. Error bars represent standard error (SE) of the mean.

Analysis of peroxidase enzymes including APx and GPx revealed consistent patterns. APx activity showed the lowest levels in control samples, with a significant increase in BV7-treated samples and the highest activity in pathogen-infected tissues (15.58±0.23 U mg^-1^ protein) (Fig 1f). GPx activity also followed mirrored this pattern, demonstrating progressive increases from control to BV7 to pathogen treatments (71.51±0.7 U mg^-1^ protein) (Fig 1g). The activities of defense enzymes showed remarkable consistency in their response patterns, with control samples consistently showing the lowest activity, BV7 treatment inducing intermediate levels, and pathogen infection triggering the highest enzymatic responses. The only exception to this pattern was observed in total protein content, where BV7 treatment induced the highest content in plant leaves. The results indicated that both beneficial (BV7) and pathogenic treatments trigger plant defensive responses, but at different intensities, with pathogenic infection generally eliciting the strongest enzymatic responses while the beneficial bacteria induces moderate levels of these defense-related enzymes. These findings therefore, suggested the need to explore the molecular basis of the interaction conditions in tomato plants at the proteomic level.

The LC-MS-based proteomic analysis revealed distinct protein expression patterns across control, beneficial bacteria (BV7)-inoculated, and pathogen-infected tomato leaves, as illustrated in the Venn diagram (Fig 2a). The analysis identified unique protein signatures for each treatment condition, with 316 proteins exclusively present in control plants, 232 proteins unique to BV7-inoculated plants, and 96 proteins specifically associated with pathogen infection. Additionally, the diagram showed overlapping proteins, with 202 proteins shared between control and BV7 inoculation, 54 proteins common between BV7 and pathogen infection, and 26 proteins shared across all the three conditions. In pathogen-infected leaves (Fig 2b), the 96 unique proteins were primarily associated with metabolic processes, cellular metabolism, and organic substance metabolic processes, as indicated by the GO term analysis. This suggested significant metabolic reprogramming in response to pathogen infection, likely representing the plant’s defense mechanisms against the pathogen. The control condition (Fig 2c) showed the highest number of unique proteins (316), with significant enrichment in cellular processes, metabolic pathways, and macromolecule metabolic processes. The presence of these proteins indicated normal physiological processes occurring in healthy tomato plants, establishing a baseline for comparison with treated conditions. In BV7-inoculated plants (Fig 2d), 232 unique proteins showed enrichment in metabolic processes, cellular processes, and various binding activities. Notably, the GO analysis revealed significant representation of proteins involved in cellular component organization, biosynthetic processes, and intracellular membrane-bounded organelle functions. This protein profile suggested that BV7 inoculation triggers specific cellular responses that may contribute to enhanced plant growth and defense preparedness.

**Figure 2.**
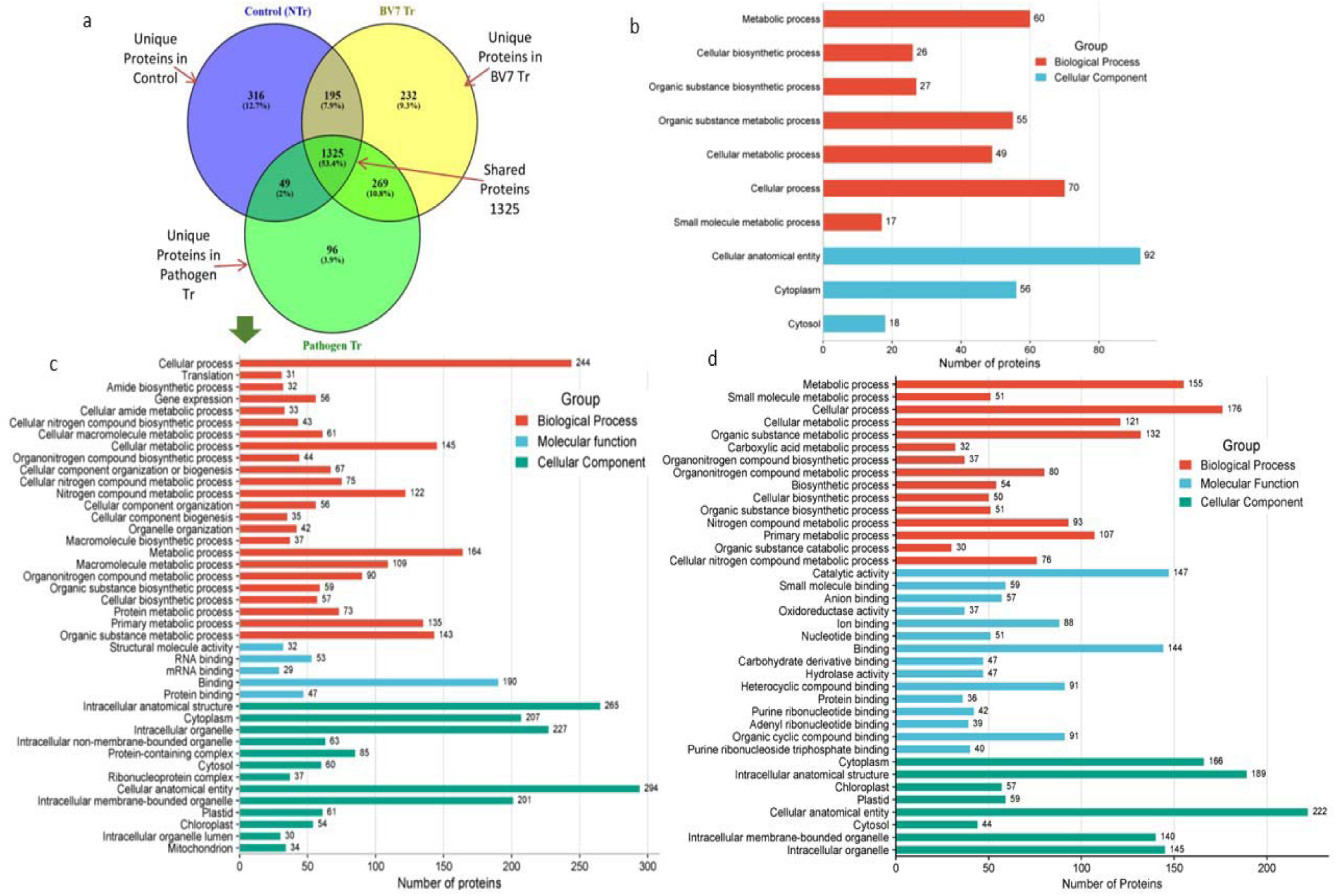
Plant leaves of tomato showing a. classification of unique proteins using venn diagram in control, BV7 beneficial bacteria inoculated and pathogen infected plant leaves as identified by LC-MS-based proteomics analysis, b. Gene Ontology (GO) terms functional classification (FC) for 96 unique proteins enriching different metabolic pathways in pathogen infected tomato plant leaves, c. FC for 316 proteins unique in control leaves, d. FC for 232 proteins unique in BV7 inoculated plant leaves. GO terms are grouped in different color bars including ‘red’ for ‘biological process’, ‘blue’ for ‘molecular function’ and ‘green’ for ‘cellular component’

A detailed analysis on how the unique proteins identified in the control, BV4 inoculated and pathogen infected plants enriched metabolic pathways, all the GO terms are presented for ‘biological process’, ‘molecular function’, ‘cellular component’, ‘subcellular localization’, ‘protein domain features’ and ‘annotated key words’ in Supplementary Tables S1, S2 and S3. STRING assigned GO terms for ‘biological process’ to the uniquely identified proteins in all the treatment conditions revealed distinct biological responses reflecting the impact of each conditions on plant health and resilience (Table 1). Majority of unique proteins in the control plants were primarily associated with cellular processes (244), cell metabolism (145), metabolic process (164), primary metabolic process (135), nitrogen compound metabolism (122), macromolecule biosynthesis (109), and organic substance metabolism (143) (Table S1). It suggested a normal plant physiological process indicating majorly protein enrichment in foundational cellular mechanism, gene expression, nitrogen metabolism, and macromolecule biosynthesis. The compartmentalization of proteins is a crucial for protein function as the specific location of proteins within cells. In control plants, unique proteins were predominantly localized to intracellular compartments, including cytoplasm, ribosomes, chloroplasts, and the mitochondria (Table S1). In BV7-inoculated plants, it was seen that besides maintaining metabolic (155), cellular (176) and cellular metabolic process (121 proteins), the biological processes for the unique proteins shifted towards enhanced small molecule biosynthesis, nitrogen metabolism, amino acid metabolism, carboxylic acid and vitamin biosynthetic processes (Table S2). Increased enrichment of proteins in vitamin biosynthesis (e.g. riboflavin) and protein refolding indicated beneficial impact of BV7 inoculation, which further caused unique protein enrichment in subcellular compartments such as the cytoplasm, chloroplasts, peroxisomes, and nucleosomes, supposedly enhancing cellular processes like metabolic activity, antioxidant defense (*via* peroxisomes), and photosynthesis (chloroplast proteins) (Table S2). In contrast, in plants infected with *A. solani,* there were unprecedented drop in the number of GO terms to which the 96 uniquely present proteins enriched (Table S3) and the enrichment showed a distinct profile dominated by catabolic and small molecule metabolic processes, indicative of stress- related responses. In pathogen-infected plants, protein localization was shown in cytoplasmic and cytosolic compartments. These proteins are involved in stress responses and metabolic disruptions, indicating a shift from constructive cellular activities to defensive mechanisms (Table S3).

**Table 1.** Gene Ontology (GO) terms for the unique proteins found in Control, BV7 inoculated and Pathogen infected tomato plants having distinguished changes in terms of the ‘biological processes’ due to interaction with *Bacillus subtilis* (BV7) and the Pathogen

| Sr. No. | GO term description | GO term ID | Observed gene count | Strength | False discovery rate (FDR) |
| --- | --- | --- | --- | --- | --- |
| 1. | Cellular process | GO:0009987 | 244 | 0.17 | 9.59E-16 |
| 2. | Translation | GO:0006412 | 31 | 0.67 | 9.66E-09 |
| 3. | Amide biosynthetic process | GO:0043604 | 32 | 0.63 | 1.41E-08 |
| 4. | Gene expression | GO:0010467 | 56 | 0.4 | 2.38E-07 |
| 5. | Cellular amide metabolic process | GO:0043603 | 33 | 0.56 | 2.38E-07 |
| 6. | Cellular nitrogen compound biosynthetic process | GO:0044271 | 43 | 0.45 | 1.13E-06 |
| 7. | Cellular macromolecule metabolic process | GO:0044260 | 61 | 0.35 | 1.36E-06 |
| 8. | Cellular metabolic process | GO:0044237 | 145 | 0.18 | 1.79E-06 |
| 9. | Organonitrogen compound biosynthetic process | GO:1901566 | 44 | 0.43 | 2.01E-06 |
| 10. | Cellular component organization or biogenesis | GO:0071840 | 67 | 0.32 | 2.39E-06 |
| 11. | Ribonucleoprotein complex biogenesis | GO:0022613 | 22 | 0.64 | 5.94E-06 |
| 12. | Cellular nitrogen compound metabolic process | GO:0034641 | 75 | 0.28 | 9.81E-06 |
| 13. | Cellular macromolecule biosynthetic process | GO:0034645 | 33 | 0.48 | 1.05E-05 |
| 14. | Ribosome biogenesis | GO:0042254 | 18 | 0.64 | 8.78E-05 |
| 15. | Nitrogen compound metabolic process | GO:0006807 | 122 | 0.18 | 0.0001 |
| 16. | Cellular component organization | GO:0016043 | 56 | 0.3 | 0.00021 |
| 17. | Cellular component biogenesis | GO:0044085 | 35 | 0.4 | 0.00021 |
| 18. | Organelle organization | GO:0006996 | 42 | 0.35 | 0.00024 |
| 19. | Macromolecule biosynthetic process | GO:0009059 | 37 | 0.37 | 0.00039 |
| 20. | Metabolic process | GO:0008152 | 164 | 0.13 | 0.00041 |
| 21. | Macromolecule metabolic process | GO:0043170 | 109 | 0.17 | 0.00063 |
| 22. | Proteasome-mediated ubiquitin-dependent protein catabolic process | GO:0043161 | 12 | 0.73 | 0.00087 |
| 23. | Organonitrogen compound metabolic process | GO:1901564 | 90 | 0.19 | 0.001 |
| 24. | Establishment of localization in cell | GO:0051649 | 23 | 0.47 | 0.0011 |
| 25. | Organic substance biosynthetic process | GO:1901576 | 59 | 0.25 | 0.0016 |
| 26. | Cellular biosynthetic process | GO:0044249 | 57 | 0.25 | 0.0018 |
| 27. | Protein metabolic process | GO:0019538 | 73 | 0.22 | 0.0019 |
| 28. | Primary metabolic process | GO:0044238 | 135 | 0.13 | 0.0026 |
| 29. | Cellular localization | GO:0051641 | 29 | 0.38 | 0.0027 |
| 30. | Generation of precursor metabolites and energy | GO:0006091 | 19 | 0.46 | 0.0076 |
| 31. | Aerobic electron transport chain | GO:0019646 | 5 | 1.09 | 0.0109 |
| 32. | Mitochondrial ATP synthesis coupled electron transport | GO:0042775 | 5 | 1.09 | 0.0109 |
| 33. | Intracellular transport | GO:0046907 | 20 | 0.43 | 0.011 |
| 34. | Organic substance metabolic process | GO:0071704 | 143 | 0.11 | 0.0126 |
| 35. | rRNA processing | GO:0006364 | 11 | 0.61 | 0.0141 |
| 36. | Ubiquitin-dependent protein catabolic process | GO:0006511 | 14 | 0.5 | 0.0239 |
| 37. | Microtubule-based process | GO:0007017 | 10 | 0.61 | 0.0251 |
| 38. | Chloroplast localization | GO:0019750 | 4 | 1.16 | 0.0251 |
| 39. | Mitochondrial electron transport, NADH to ubiquinone | GO:0006120 | 3 | 1.37 | 0.0413 |
| 40. | Electron transport chain | GO:0022900 | 12 | 0.51 | 0.0414 |
| 41. | Proteolysis involved in protein catabolic process | GO:0051603 | 15 | 0.44 | 0.0476 |

b. GO enrichment terms (41) for the 'biological process' of unique proteins in BV7 inoculated plants
| Sr. No. | GO term description | GO term ID | Observed gene count | Strength | False discovery rate (FDR) |
| --- | --- | --- | --- | --- | --- |
| 1. | Metabolic process | GO:0008152 | 155 | 0.24 | 4.33E-14 |
| 2. | Small molecule metabolic process | GO:0044281 | 51 | 0.58 | 6.27E-13 |
| 3. | Cellular process | GO:0009987 | 176 | 0.16 | 4.70E-10 |
| 4. | Cellular metabolic process | GO:0044237 | 121 | 0.24 | 3.70E-09 |
| 5. | Organic substance metabolic process | GO:0071704 | 132 | 0.21 | 8.17E-09 |
| 6. | Carboxylic acid metabolic process | GO:0019752 | 32 | 0.62 | 1.17E-08 |
| 7. | Cellular amino acid metabolic process | GO:0006520 | 21 | 0.77 | 8.86E-08 |
| 8. | Small molecule biosynthetic process | GO:0044283 | 24 | 0.68 | 1.88E-07 |
| 9. | Organonitrogen compound biosynthetic process | GO:1901566 | 37 | 0.49 | 7.80E-07 |
| 10. | Organonitrogen compound metabolic process | GO:1901564 | 80 | 0.28 | 1.15E-06 |
| 11. | Biosynthetic process | GO:0009058 | 54 | 0.32 | 3.53E-05 |
| 12. | Cellular amino acid biosynthetic process | GO:0008652 | 12 | 0.87 | 5.31E-05 |
| 13. | Cellular biosynthetic process | GO:0044249 | 50 | 0.33 | 5.88E-05 |
| 14. | Carboxylic acid biosynthetic process | GO:0046394 | 17 | 0.67 | 6.47E-05 |
| 15. | Organic substance biosynthetic process | GO:1901576 | 51 | 0.32 | 7.14E-05 |
| 16. | Catabolic process | GO:0009056 | 34 | 0.42 | 8.83E-05 |
| 17. | Alpha-amino acid metabolic process | GO:1901605 | 14 | 0.74 | 0.00011 |
| 18. | Nitrogen compound metabolic process | GO:0006807 | 93 | 0.19 | 0.00028 |
| 19. | Alpha-amino acid biosynthetic process | GO:1901607 | 10 | 0.87 | 0.00036 |
| 20. | Primary metabolic process | GO:0044238 | 107 | 0.17 | 0.00043 |
| 21. | Organic substance catabolic process | GO:1901575 | 30 | 0.41 | 0.00043 |
| 22. | Organonitrogen compound catabolic process | GO:1901565 | 19 | 0.5 | 0.0027 |
| 23. | Nucleosome assembly | GO:0006334 | 5 | 1.2 | 0.0042 |
| 24. | Cellular catabolic process | GO:0044248 | 24 | 0.41 | 0.005 |
| 25. | Protein refolding | GO:0042026 | 5 | 1.12 | 0.0081 |
| 26. | Cellular nitrogen compound metabolic process | GO:0034641 | 50 | 0.24 | 0.0115 |
| 27. | Cellular nitrogen compound biosynthetic process | GO:0044271 | 26 | 0.36 | 0.0115 |
| 28. | Hydrocarbon biosynthetic process | GO:0120251 | 5 | 1.04 | 0.0174 |
| 29. | Generation of precursor metabolites and energy | GO:0006091 | 15 | 0.49 | 0.0201 |
| 30. | Olefinic compound biosynthetic process | GO:0120255 | 5 | 1 | 0.0252 |
| 31. | Riboflavin biosynthetic process | GO:0009231 | 3 | 1.47 | 0.027 |
| 32. | Peptide metabolic process | GO:0006518 | 16 | 0.45 | 0.0291 |
| 33. | Vitamin biosynthetic process | GO:0009110 | 5 | 0.94 | 0.0333 |
| 34. | Ethylene biosynthetic process | GO:0009693 | 4 | 1.13 | 0.0333 |
| 35. | Pigment metabolic process | GO:0042440 | 6 | 0.83 | 0.0333 |
| 36. | Protein-containing complex assembly | GO:0065003 | 13 | 0.5 | 0.0333 |
| 37. | Organophosphate metabolic process | GO:0019637 | 15 | 0.45 | 0.0338 |
| 38. | Cellular amide metabolic process | GO:0043603 | 17 | 0.41 | 0.0374 |
| 39. | NADP metabolic process | GO:0006739 | 4 | 1.08 | 0.0384 |
| 40. | Monocarboxylic acid metabolic process | GO:0032787 | 12 | 0.5 | 0.0443 |
| 41. | Chromatin assembly | GO:0031497 | 6 | 0.78 | 0.0477 |

c. GO enrichment terms (7) for the 'biological process' of unique proteins in Pathogen inoculated plants
| Sr. No. | GO term description | GO term ID | Observed gene count | Strength | False discovery rate (FDR) |
| --- | --- | --- | --- | --- | --- |
| 1. | Metabolic process | GO:0008152 | 60 | 0.21 | 0.0055 |
| 2. | Cellular biosynthetic process | GO:0044249 | 26 | 0.43 | 0.0055 |
| 3. | Organic substance biosynthetic process | GO:1901576 | 27 | 0.43 | 0.0055 |
| 4. | Organic substance metabolic process | GO:0071704 | 55 | 0.21 | 0.0059 |
| 5. | Cellular metabolic process | GO:0044237 | 49 | 0.23 | 0.012 |
| 6. | Cellular process | GO:0009987 | 70 | 0.14 | 0.0242 |
| 7. | Small molecule metabolic process | GO:0044281 | 17 | 0.48 | 0.0242 |

The proteomes of two plant groups involving control *versus* BV7-inoculated (C*vs*BV7) and control *versus* pathogen infected (C*vs*Pathogen) plants were mutually compared to understand changes at the level of protein profiles. Univariate analysis using volcano plot indicated regulation (up- or -down) of proteins in both the plant groups in terms of fold change (FC>1.5) and *p* values (<0.05) (Fig 3a,b). C*vs*BV7 plant group showed 101 differentially up-regulated and 106 down-regulated proteins (Fig 3a). In C*vs*Pathogen plant group, there were 75 up- and 108 down-regulated proteins (Fig. 3b). The heatmap generated a correlation between overall proteins in the C*vs*BV7 (Fig. 3c) and in C*vs*Pathogen group (Fig 3d). Heatmap for CvsBV7 showed a clear segregation into four distinct clusters (represented by the red and blue blocks). The sharp contrast between these blocks indicated that BV7 treatment leads to consistent and substantial changes in the overall protein composition compared to control conditions. The presence of both positive (red) and negative (blue) correlations suggested a complex protein regulation networks are activated during beneficial bacterial colonization. The heatmap for CvsPathogen displayed a more complex correlation pattern with multiple smaller clusters, indicating that pathogen infection triggers a more diverse and perhaps more specialized protein response. The heatmap showed more gradual transitions between correlation clusters, suggesting interconnected protein regulation networks responding to pathogen stress. The pattern further differentiated the samples based on protein composition and indicated major changes in the overall protein profile in BV7 inoculated and pathogen infected plants, indicating distinct changes at the protein regulation level in tomato in response to both the beneficial and pathogen interaction.

**Figure 3.**
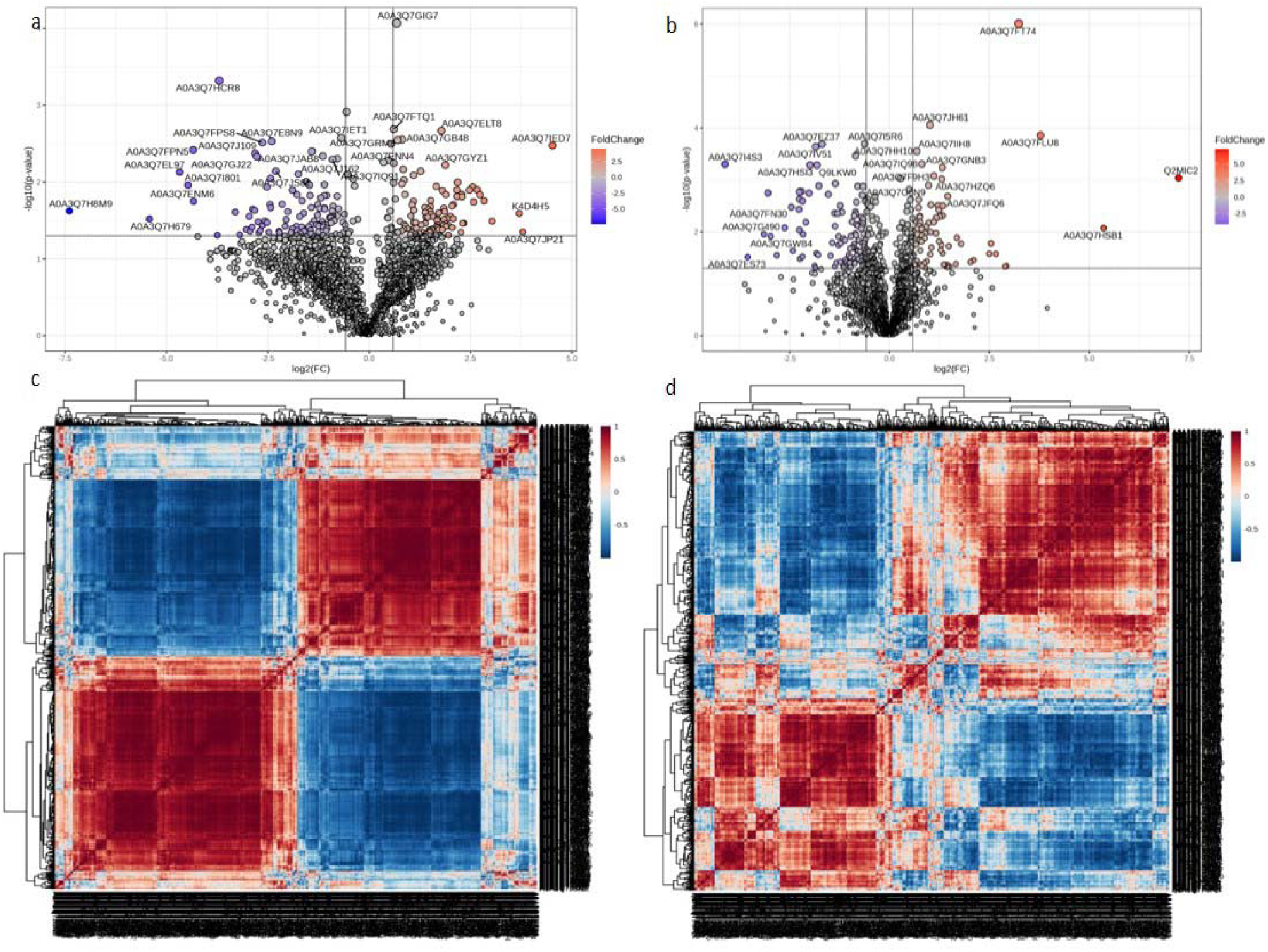
Identification of proteins that differ significantly in terms of their fold change (FC) values at p<0.05 to describe protein regulation (up- and down) in the samples, Volcano plot for a. CvsBV7 and b. CvsPathogen plant group; and correlation heatmap showing sample differentiation based on overall protein composition in c. CvsBV7 and d. CvsPathogen plant group

A distinct but complex enrichment of unique proteins identified in C*vs*BV7 (red lines) and C*vs*Pathogen (blue lines) plant groups in various metabolic pathways was observed in tomato leaves in the pathway map generated by iPath3.0 highlighting unique protein engagement patterns (Fig. 4). Contrast biosynthetic and metabolic pathways were mapped in iPATH by unique proteins found in the plant group Cvs BV7 and CvsPathogen. In CvsBV7-inoculated plant group, there is notable activation of pathways related to primary metabolism, including enhanced photosynthetic processes and energy metabolism. BV7-treated plants show enhanced protein engagement in pathways related to beneficial metabolite production, suggesting improved plant fitness. The positive influence of BV7 interaction on the plant’s energy production and carbon fixation capabilities is supposed to potentially contribute to improved plant growth and development. The unique proteins in Cvspathogen-infected plants showed differential engagement in stress response pathways and secondary metabolism. Inoculation of beneficial bacteria BV7 enriched vitamin metabolism including that of thiamine, riboflavin, and folate metabolism more prevalently as compared to the unique proteins in CvsPathogen plant group. Likewise, phenylpropanoid biosynthesis, tyrosine and phenylalanine metabolism was highly enriched by the proteins unique in pathogen infected plants leaves showing significant activation of defense-related pathways, including those involved in oxidative stress response. This has indicated plant’s shift towards defense mechanisms rather than growth-promoting processes under pathogenic interaction condition.

**Figure 4.**
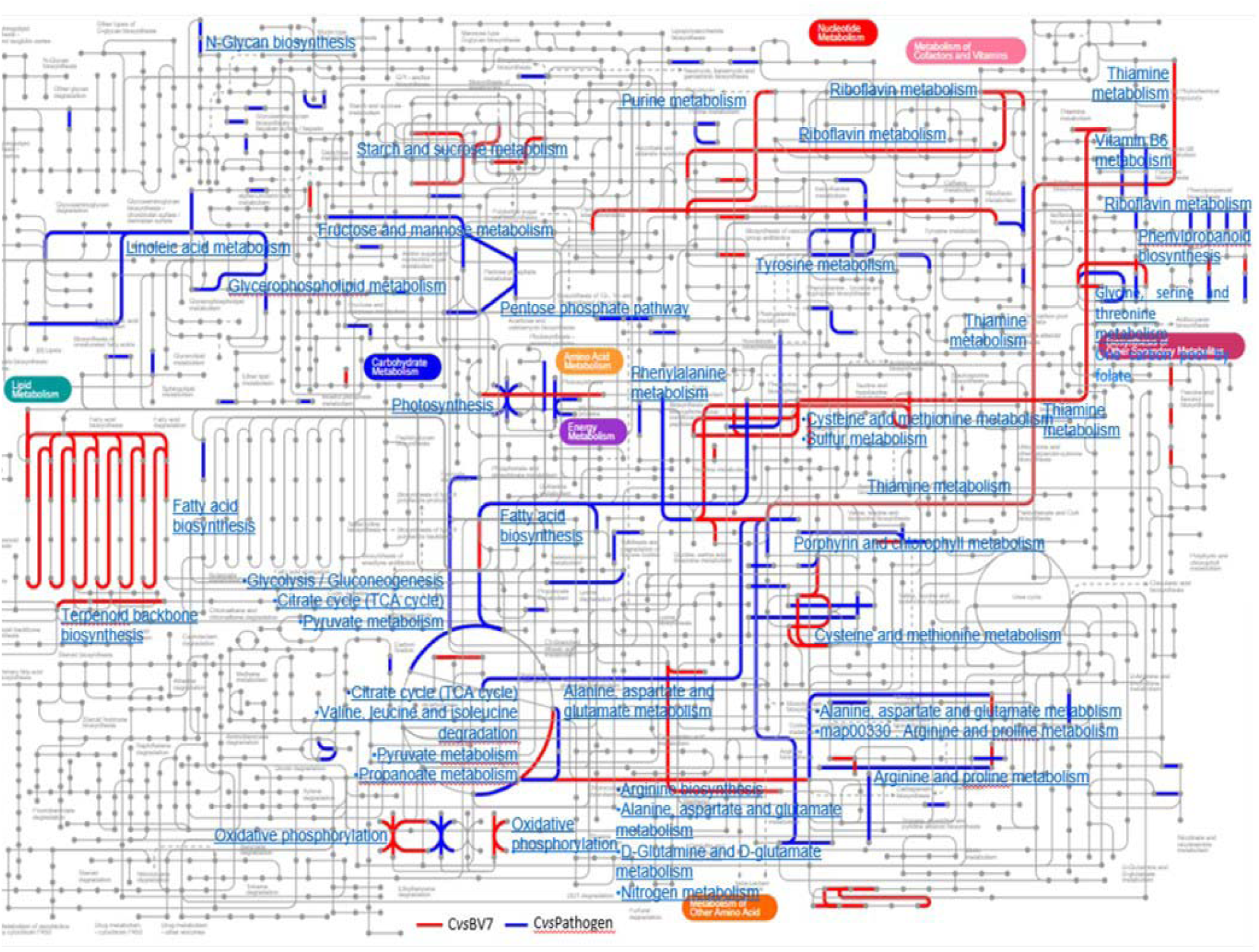
Comparative pathway analysis of differentially expressed proteins in plants under different treatments. The metabolic pathway map illustrates unique proteins identified in BV7- inoculated plants (red lines, CvsBV7) and pathogen-infected plants (blue lines, CvsPathogen) relative to control plants. The network demonstrates the complex interconnections between various metabolic pathways (shown in blue text), including but not limited to photosynthesis, fatty acid metabolism, amino acid metabolism, and secondary metabolite biosynthesis. The intricate pathway mapping reveals distinct metabolic responses triggered by beneficial (BV7) and pathogenic interactions in plants, highlighting the differential regulation of key biological processes under these conditions.

Particularly noteworthy is the differential regulation of amino acid metabolism and secondary metabolite biosynthesis between the two conditions. The interconnectedness of these pathways demonstrated complex metabolic reprogramming during plant-microbe interactions. The beneficial bacterium BV7 appeared to promote pathways that enhance plant growth and development, while pathogen infection triggered extensive metabolic changes related to stress response and defense. This differential pathway engagement has provided insights into how plants modulate their metabolism in response to beneficial versus pathogenic interactions and how the treatments underline plant’s ability to distinguish between beneficial and harmful microorganisms, leading to appropriate metabolic responses that either promote growth or activate defense mechanisms.

Identification of reliable biomarkers has remained central among the major aims of any proteomic analysis (Alharbi et al, 2020). Results based on ROC analysis identified potential protein biomarkers distinguishing between C*vs*BV7 inoculation (Fig 5a) and C*vs*Pathogen (Fig 5b). The comparative analysis of biomarker proteins revealed distinct molecular signatures in plants responding to beneficial bacteria (BV7) versus pathogen infection. In BV7-inoculated plants (Fig. 5a), PSI-K (Photosystem I subunit K) emerged as the most significant biomarker with the highest average importance score (2.53), followed by ribosomal protein L35, Glycine-rich protein, Alcohol acyl transferase, alanine transaminase, Glycine-rich protein, Fruit-ripening protein, Glucan endo-1,3-beta-D-glucosidase. The predominance of photosynthesis-related and metabolic proteins among the top biomarkers suggested that BV7 inoculation enhances primary metabolism and energy production processes. Notably, several stress-response proteins, including UTP-glucose-1-phosphate uridylyltransferase and NADP-binding domain-containing proteins, also showed high importance scores (>1.8), indicating an induced balanced regulation of defense responses along with the growth and development. In contrast, pathogen-infected plants (Fig. 5b) displayed a distinct set of biomarker proteins primarily associated with defense responses. The identified biomarker proteins Cytochrome b559 subunit alpha ranked highest (2.38), followed by the proteins ethylene-responsive proteinase inhibitor 1, thiamine thiazole synthase, glutamate synthase (ferredoxin), carboxypeptidase A inhibitor-like domain-containing protein, and 14-3-3 protein 7 in CvsPathogen plant group (Fig. 5b) indicated the protein profile shift towards signal, pathogenesis-related stress response, catabolic processes and defense mechanisms during pathogen infection. The functional distribution of proteins in different groups viz. protein synthesis and ribosomal proteins, metabolic enzymes, proteins related to plant growth, photosynthesis, chloroplast, cell wall modification, energy metabolism, protein modification and regulation, stress response, defense and signal transduction highlighted the sophisticated molecular machinery plants employ to distinguish between beneficial and pathogenic interactions, appropriately allocating resources either toward growth promotion or defense responses. The differential regulation of these protein groups demonstrated the plant’s ability to mount specific and appropriate responses to different microbial encounters.

**Figure 5.**
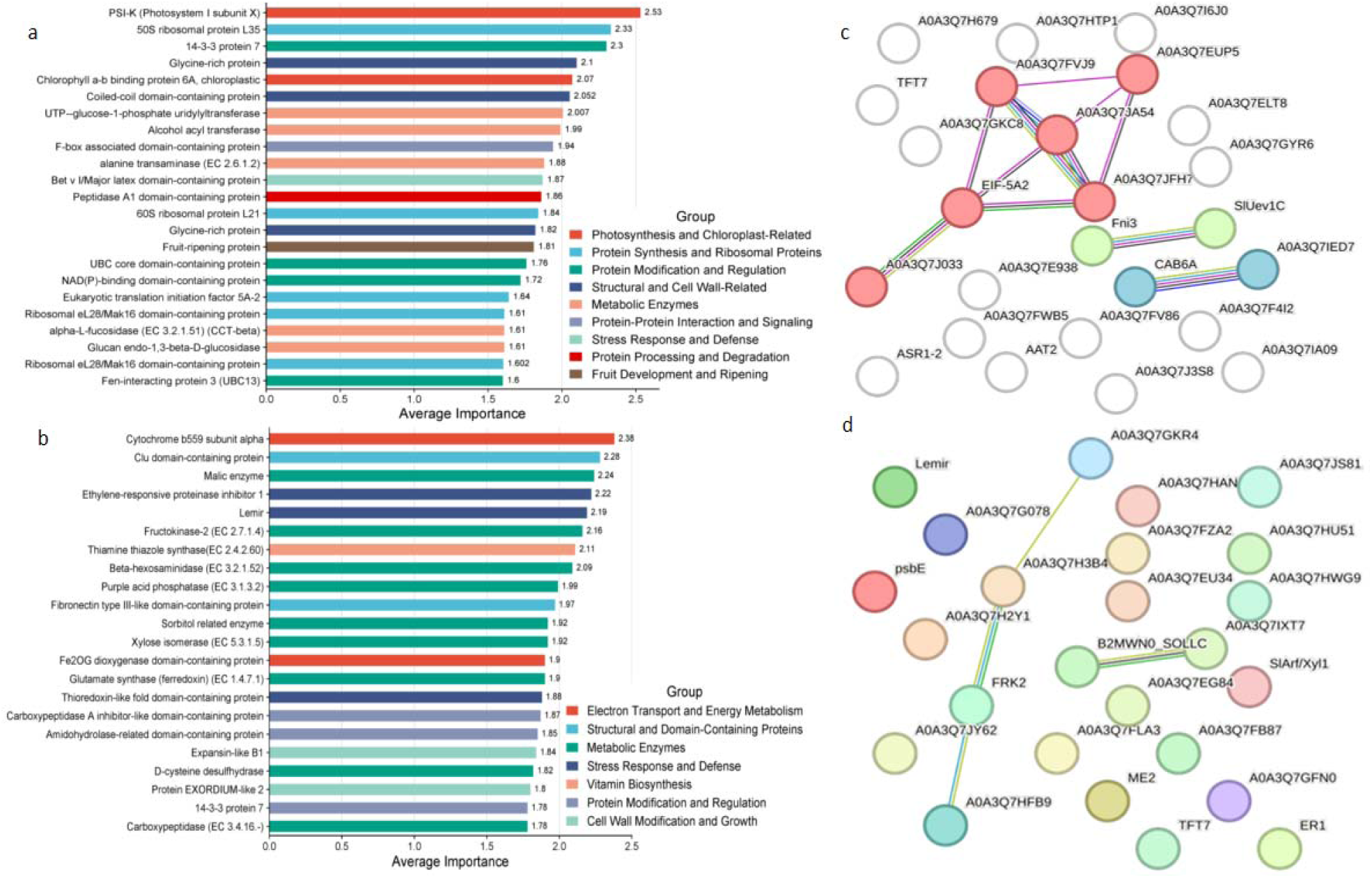
Identification of biomarker proteins (using ROC curve analysis classification method PLS-DA and feature ranking method PLS-DA built-in in MetaboAnalyst 6.0) and their interactions using String database in CvsBV7 and CvsPathogen plant groups. Top 25 biomarker proteins ranked by average importance along with their functional groups were identified in a. BV7-inoculated and b. pathogen infected plants compared to control. Red and green bars indicate the average importance scores of proteins in CvsBV7 and CvsPathogen plant groups, respectively; c. Protein-protein interaction (PPI) network of key biomarker proteins in CvsBV7 group, with nodes colored according to functional clusters and edge thickness representing interaction strength, d. PPI in CvsPathogen plant group showing interactions and functional clustering representing differential molecular responses to beneficial and pathogenic interactions

The protein-protein interaction networks provided further insights into the functional organization of these responses. In BV7-treated plants (Fig. 5c), the network showed concentrated clustering around metabolic processes, with strong interactions between key proteins (indicated by edge thickness). The functional clusters, represented by differently colored nodes, suggest coordinated regulation of growth-promoting pathways. The pathogen-response network (Fig. 5d) exhibited a more dispersed interaction pattern with multiple functional clusters, reflecting the complex nature of defense responses. This network organization demonstrates the activation of diverse defense mechanisms and stress response pathways during pathogen infection. These findings highlighted how plants engage different protein networks and metabolic pathways when interacting with beneficial versus pathogenic microorganisms, providing molecular insights into the plant’s ability to respond to different microbial interactions. Overall, our results have pointed out the complex and specific protein-level responses of plants to varied microbial interactions, providing valuable insights into the molecular mechanisms underlying plant-microbe interactions.

## 4. Discussion

The study evaluated proteome level biochemical dynamic changes in the leaves of tomato (var. Kashi Aman) plants inoculated with *Bacillus* BV7 (beneficial) and infected with *A. solani* (pathogenic) interaction using comparative proteomics. The findings revealed critical insights into how beneficial and pathogenic interactions shape the plant’s proteome, influencing its metabolic, biosynthetic, and defense-related processes. Both the beneficial bacterial inoculation and pathogen infection in tomato affected defense related antioxidant enzyme activity and presented proteome level distinct changes that reflected molecular mechanisms underlying plant-microbe interactions. An exclusive data analysis pipeline, including proteome extraction, data generation, enzyme assay, protein annotation, pathway enrichment analyses and statistical models was employed to identify significantly regulated proteins and biomarkers. Mapped altered pathways and associated biological processes were implied to generate a comprehensive view of the cellular reprogramming events in the plants (Khan et al, 2024).

The antioxidant enzymes such as POD, SOD, APx, and GPx play pivotal roles in plant defense by mitigating oxidative stress caused by biotic interactions (Rajput et al, 2021). Significant increase in enzyme activities observed in BV7-inoculated plants highlighted their role in priming plant defense mechanisms. Significantly higher protein content in BV7-treated plants aligns with findings by Hashem et al (2019), who demonstrated that beneficial *Bacillus* strains enhance protein synthesis machinery in host plants. The lower protein content in pathogen-infected tissues may indicate protein degradation or metabolic disruption during pathogenic invasion, consistent with observations by Schmey et al. (2024). PAL activity showed progressive elevation from control to BV7 to pathogen treatments, with pathogen infection inducing the highest activity. This pattern is of particular significance as the enzyme PAL represents the first committed step in the phenylpropanoid pathway, crucial for defense compound synthesis (Vogt 2010). The moderate induction of PAL enzyme activity by the *Bacillus* BV7 inoculation suggested a balanced immune response that may have contributed to improved plant health (Bhattacharyya et al 2020).

The activity of SOD was highest in pathogen-infected tissues indicating intense oxidative stress management level, as previously noted by Torres et al (2006) during plant-pathogen interactions (Kunos et al 2022). The moderate SOD induction in *Bacillus* BV7-treated plants suggested a primed state of oxidative stress response without the metabolic burden of full activation, supporting the concept of beneficial microbe-induced systemic resistance described by Pieterse et al. (2014). The elevated GPx activity in pathogen-infected plants reflected deployment of ROS-scavenging mechanisms, crucial for managing oxidative stress during pathogen attack (Wang et al 2019). The intermediate levels of peroxidase activities in BV7-treated plants have led to presume a calibrated defense response that may contribute to enhanced stress tolerance without compromising growth in beneficial microbe-plant interactions (Meena et al. 2017, Tharanath et al, 2024).

The comparative proteomic analysis has revealed distinctive proteomic responses in tomato under different microbial interactions, highlighting the proteome-level mechanisms plants employ to differentiate between beneficial and pathogenic interactions. In control plants, presence of 316 unique proteins enriched in fundamental cellular and metabolic processes represented a robust baseline of physiological activities. This protein profile indicated an active primary metabolism regulating photosynthesis and protein synthesis and the enzymes essential for normal plant growth and development (Cheng et al, 2010, Tcherkez et al, 2020, Ji et al, 2022). The beneficial bacteria (BV7) inoculation induced a distinct protein regulation pattern with 232 unique proteins, demonstrating proteins related to photosynthesis, primary metabolism, vitamin biosynthesis (particularly riboflavin), and protein refolding mechanisms. The activation of these growth-promoting pathways, coupled with maintained basal defense readiness, has shown that beneficial microbial interaction optimally regulated plant development while ensuring protective mechanisms to remain functional (Mhlongo et al 2020). In contrast, pathogen-infected plants exhibited a more focused response with 96 unique proteins, predominantly associated with stress response and defense-related enzymes. The increased catabolic processes suggested significant metabolic reprogramming towards defense at the expense of growth (Bentham et al, 2020). This protein level shift represented a resource allocation strategy where plants prioritize survival over growth under pathogenic stress (Huot et al 2014). The protein-protein interaction networks further demonstrated differential responses (Hollander et al 2021). BV7-induced networks showed concentrated clustering with strong metabolic interactions, indicating systematic enhancement of growth and development processes. Conversely, pathogen-induced networks displayed more dispersed clustering with multiple functional groups, reflecting the activation of diverse stress response mechanisms (Zhang et al., 2022). Interestingly, the common elements of 54 shared proteins in beneficial and pathogenic response pathways, potentially indicated basal defense priming by beneficial bacteria (Mhlongo et al., 2018).

The results of volcano plot analysis effectively illustrated the magnitude and significance of protein regulation in both the plant groups. In the CvsBV7 comparison, relatively balanced distribution of up- and down-regulated proteins (fold change >1.5, p<0.05) suggested a coordinated metabolic reprogramming in response to beneficial bacterial colonization (Li et al, 2023). The comparable numbers of proteins regulated in both directions indicated that the inoculation of *Bacillus* BV7 induced both activation and suppression of specific cellular processes rather than a unidirectional response (Salehin et al, 2021). In contrast, the CvsPathogen group exhibited higher number of down-regulated proteins suggesting that pathogen infection has caused broader suppression of normal cellular processes, possibly reflecting the pathogen’s strategy to compromise host defense mechanisms (Huot et al, 2014).

The heatmap of CvsBV7 demonstrated clear segregation into four distinct clusters indicating that *Bacillus* BV7 treatment leads to well-defined and consistent changes in protein composition compared to control condition. The presence of both positive and negative correlations suggested the activation of complex protein regulation networks during beneficial bacterial colonization, likely reflecting the simultaneous promotion of growth and basal defense mechanisms (Karasov et al, 2017). The heatmap of CvsPathogen revealed a complex correlation with multiple smaller clusters. This intricate organization suggested that pathogen infection triggered more diverse and specialized protein response compared to beneficial bacterial interaction. The gradual transitions between correlation clusters indicated interconnected protein regulation networks responding to pathogen stress, highlighting the complexity of defense responses (Ding et al., 2024). Distinct patterns observed in both heatmaps effectively differentiated the samples based on protein composition, indicating major changes in the overall protein profile in both BV7-inoculated and pathogen-infected plants. This differentiation at the protein level suggested proteome-based distinguished machinery in plants for beneficial and pathogenic interactions, leading to appropriate physiological responses (Hu et al., 2015).

The metabolic pathway maps by iPath 3.0 revealed that BV7 treatment activated pathways related to enhanced photosynthesis, primary metabolism, energy production, and vitamin synthesis, while pathogen infection triggered pathways associated with stress response, defense mechanisms, and secondary metabolism (Kong et al., 2023). The enhanced engagement of photosynthetic processes and energy metabolism pathways in BV7-inoculated plants suggested that beneficial bacteria promoted plant’s energy and metabolic efficiency (Stringlis et al., 2018). The BV7 interaction showed enhanced activity in carbon fixation and energy production pathways that potentially contribute to improved plant growth and development (Tiwari et al, 2022). Such observations correlate with the positive influence of beneficial bacteria on plant fitness and productivity (Harman et al, 2021). The differential pathway engagement demonstrated plant’s modulation of their metabolism in response to different microbial interactions (Nishad et al., 2020).

Particularly striking is the activation of pathways related to beneficial vitamin metabolism pathways. Increased engagement of proteins in thiamine, riboflavin, and folate metabolism pathways during BV7 interaction compared to pathogen-infected plants suggested positive influence of the beneficial interaction on plant’s nutritional and metabolic status (Vassilev and Malusà, 2021). Contrary to this, distinct metabolic profile dominated by pathways involved in stress response and secondary metabolism was observed in the CvsPathogen group. The enhanced engagement of proteins in phenylpropanoid biosynthesis, tyrosine, and phenylalanine metabolism pathways indicated significant activation of defense-related mechanisms. This metabolic shift is presumed to be due to the plant’s reallocation of resources from growth-promoting processes to defense mechanisms under pathogenic stress (Schultz et al., 2013). Differential regulation of amino acid metabolism and secondary metabolite biosynthesis between the two conditions is particularly noteworthy. The interconnectedness of these pathways, as visualized in the network map, demonstrates the complex metabolic reprogramming that occurs during plant-microbe interactions. The beneficial bacterium BV7 has appeared to promote pathways that enhance plant growth and development, while pathogen infection triggers extensive metabolic changes related to stress response and defense (Shan et al, 2024).

The analysis of protein biomarkers revealed crucial insights into the distinctive molecular signatures characterizing plant responses to beneficial bacterial inoculation (BV7) and pathogen infection. In BV7-inoculated plants, PSI-K (Photosystem I subunit K) emerged as the most significant biomarker, followed by key proteins ribosomal protein L35, glycine-rich protein, alcohol acyl transferase, and alanine transaminase. The identified biomarker proteins were majorly linked to photosynthetic efficiency and energy production, protein synthesis and cellular growth, light harvesting and photosynthesis, carbohydrate metabolism, cell wall modification and fruit development processes. The predominance of photosynthesis-related and metabolic proteins among the biomarkers suggested activated primary metabolism and energy production processes (Stringlis et al., 2018). Several stress-response proteins, including UTP-glucose-1- phosphate uridylyltransferase and NADP-binding domain-containing proteins, showed high importance scores (>1.8), indicating that beneficial bacteria help maintain a balanced regulation between growth promotion and defense preparedness (Timofeeva et al., 2024).

In contrast, pathogen-infected plants showed a distinct set of biomarker proteins primarily associated with plant defense and stress response. The cytochrome b559 subunit alpha is associated with photosynthetic electron transport and ROS signaling during stress (Didaran et al, 2024), ethylene-responsive proteinase inhibitor 1+ integrates signaling between ethylene and jasmonate pathway for plant defense (Lorenzo et al, 2003), thiamine thiazole synthase interacts with abscisic acid (ABA) signaling (Ki et al, 2016), 14-3-3 protein 7 is associated with immunity-linked programmed cell death pathway in tomato (Oh et al 2010), expansin-like B1 protein regulate developmental processes (Sun et al, 2021) and thioredoxin-like proteins mediate redox signaling in plant immunity (Jedelská et al, 2020). Dominance of such proteins upon pathogenic interaction indicates a shift towards stress response, catabolic processes, and defense mechanisms during infection (Jones et al., 2019).

Protein-protein interaction (PPI) networks provided further insights into the functional organization of these responses. In BV7-treated plants, the network exhibits concentrated clustering around metabolic processes, with strong interactions between key proteins as indicated by edge thickness. The functional clusters, represented by differently colored nodes, suggest coordinated regulation of growth-promoting pathways. The pathogen-response network showed a more dispersed interaction pattern with multiple functional clusters, reflecting the complex nature of defense responses and indicating the activation of diverse defense mechanisms.

Functional distribution of proteins across different groups (protein synthesis and ribosomal proteins, metabolic enzymes, proteins related to plant growth, etc.) highlighted the sophisticated molecular machinery plants employ to distinguish between beneficial and pathogenic interactions. This differential regulation demonstrated the plant’s ability to mount specific and appropriate responses to different microbial encounters.

## 5. Conclusion

This comprehensive proteomics-based investigation has provided insights into how plants orchestrate distinct molecular responses to beneficial and pathogenic interactions. The study demonstrated that beneficial bacteria (BV7) induces a balanced protein expression profile that promotes growth while maintaining defense readiness, evidenced by 232 unique proteins predominantly involved in primary metabolism and energy production. In contrast, pathogen infection has triggered a more focused defensive response with 96 unique proteins primarily associated with stress response and defense mechanisms. The identification of specific biomarker proteins - PSI-K (score 2.53) for beneficial interactions and Cytochrome b559 (score 2.38) for pathogenic responses provided valuable molecular signatures for distinguishing between different plant-microbe interactions. The identification of 54 shared proteins between BV7 and pathogen treatments suggested a common molecular framework that may contribute to induced systemic resistance, highlighting the sophisticated nature of plant immune responses. Particularly noteworthy is the differential regulation of metabolic pathways, which uniquely revealed that beneficial bacteria (BV7) has enriched vitamin metabolism (thiamine, riboflavin, folate) and primary metabolic processes while pathogen infection activated defense-related pathways including phenylpropanoid biosynthesis, amino acid metabolism and stress responses. This has clearly demonstrated distinct metabolic reprogramming strategies in plants responding to different microbial interactions that distinguish between beneficial and harmful microorganisms. The findings have significant implications for agricultural applications, particularly in developing strategies for enhancing crop productivity and disease resistance. The identified protein biomarkers and metabolic pathways could serve as potential targets for improving plant health and developing more effective biological control methods. Future work building on these insights could lead to innovative approaches in sustainable agriculture and plant protection strategies.

## Notes

### Competing Interest Statement

The authors have declared no competing interest.

### Summary of Updates

Supplementary Tables S1, S2 and S3 are updated.

